# Microclimate habitats of *Culex* mosquitoes in urban desert environments

**DOI:** 10.64898/2025.12.12.694062

**Authors:** Chloe Martz, Alex Vela, Kelsey Lyberger

**Affiliations:** College of Integrative Sciences and Arts, Arizona State University, Mesa, AZ

**Keywords:** thermal tolerance, climate, microhabitat, urban heat, vector ecology

## Abstract

*Culex* mosquitoes are important vectors of West Nile virus, and, as small-bodied ectotherms, are sensitive to temperature. This is especially important for juvenile stages which are confined to the body of water where they hatched and are unable to relocate for thermoregulation. In an urban desert like Maricopa County, Arizona, the macroclimate is strongly influenced by the urban heat island effect where larval thermal maxima are frequently exceeded. Understanding the local temperatures (microclimates) of sites that mosquitoes chose for egg-laying and larval development is important for managing mosquito populations and reducing disease risk. We deployed small temperature loggers at multiple breeding sites throughout the city and compared those temperatures to gridded reanalysis macroclimate data. We found that the recorded aquatic microclimates were consistently cooler than corresponding macroclimates, and that higher canopy cover further buffered hot temperatures. This suggests that using interpolated macroclimate data to estimate mosquito population dynamics overestimates the thermal stress local mosquitoes actually experience. By using microclimate measures we can better predict mosquito survival and development, allowing us to better understand disease risk both now and with future warming.

## Introduction

Because insects are ectothermic, nearly all aspects of their performance, from metabolism and growth to behavior and reproduction, depend on external temperatures (Kingsolver & Huey 2008). This physiological dependence makes insects particularly sensitive to environmental variability, where temperature has long been recognized as one of the most important abiotic factors governing their abundance, phenology, and distribution (Forister and Shapiro 2003; Angilletta 2009; Sunday et al. 2019). Subtle shifts in thermal conditions can alter trophic interactions and reorganize entire insect communities (Dell et al. 2014; Haase et al. 2019). For mosquitoes, which develop in thermally dynamic aquatic habitats and as adults encounter distinct thermal challenges in the terrestrial environment, these sensitivities are pronounced and influence the transmission dynamics of vector-borne diseases (Mordecai et al. 2019). For larvae in particular, they are confined to discrete aquatic habitats that offer little opportunity to escape unfavorable temperatures. Even minor differences in local thermal regimes can accelerate or suppress development (Trudgill et al. 2005; Kingsolver and Buckley 2020), alter adult body size and fecundity (Berger et al. 2008; Tseng et al. 2018; Lyberger et al. 2024), and change the rate at which pathogens replicate within their bodies (Samuel et al. 2016; Bellone & Failloux 2020).

Among mosquito vectors, *Culex* species play a particularly important role in public health because they transmit West Nile virus (WNV) across much of North America (Sardelis et al. 2001). WNV is responsible for the majority of mosquito-borne disease cases reported annually in the United States (CDC, 2024), and transmission can lead to large outbreaks, including the largest county level outbreak recorded in the country, which occurred in Maricopa County, Arizona in 2021 *(Kretschmer et al. 2023).* Two of the most widespread species, *Culex quinquefasciatus* and *Culex tarsalis*, are considered the primary vectors of WNV in the southwest and western regions of the United States (Andreadis 2012). The range of *Cx. quinquefasciatus* extends across the southern half of the country, whereas the range of *Cx. tarsalis* extends across much of the country and northward into southern Canada (Gorris et al. 2021). They span an extraordinary range of climatic conditions including humid subtropical regions, temperate forests, and hot deserts. Understanding how they maintain viable populations in such environments is essential for predicting when and where WNV transmission risk will occur, especially as global temperatures rise and urban heat intensifies.

Urban desert environments represent some of the most thermally challenging habitats for mosquitoes. Among global mosquito habitats, desert biomes have the smallest estimated thermal safety margins across several vector species (Couper et al. 2024). The low humidity and limited precipitation characteristic of these regions further add physiological stress (Holmes & Benoit 2019). Additionally, in cities, urban heat islands can elevate local temperatures several degrees above surrounding rural areas (Oke 1982; LaDeau et al. 2015). Maricopa County in Arizona, which includes Phoenix, is one such environment where summer air temperatures routinely exceed 43°C (National Weather Service), often surpassing the upper thermal limits for *Culex* larval survival reported in laboratory studies (Mordecai et al. 2019). Yet *Culex* mosquitoes remain abundant and widespread throughout the metropolitan area, reproducing successfully through the hottest months (Wilke et al. 2023). This persistence suggests that thermal refuges at the microscale play a critical role in sustaining populations under otherwise lethal macroclimatic conditions. Small pools shaded by vegetation or infrastructure may remain several degrees cooler than surrounding air, providing a buffer that allows larvae to survive and develop. However, because these fine-scale habitats are highly localized and difficult to capture in regional datasets, most predictive models rely on coarse weather-station or reanalysis data that overlook this buffering. This has left the magnitude and ecological significance of microclimate effects poorly quantified at these hot extremes.

In this study, we characterize the microclimates of *Culex tarsalis* and *Culex quinquefasciatus* larval habitats in an urban desert and evaluate how these conditions differ from the macroclimate data typically used in ecological models. Because these species persist through extreme summer heat, quantifying the thermal environment they actually experience is essential for understanding the conditions that support their survival. By comparing microclimate temperature data from a variety of larval sites across Maricopa County to macroclimate data, we seek to answer the following: (1) Are *Culex* larval sites significantly cooler than macroclimate data during periods of peak thermal stress? (2) Do temperatures recorded at *Culex* larval sites remain within the bounds of larval thermal tolerance? (3) Does canopy cover represent a significant factor in reducing temperature in larval microhabitats? Together, these results will support more reliable predictions of where and when *Culex* populations can develop under increasingly extreme thermal conditions.

## Methods

### Study Sites and Sample Collection

Field sampling was conducted in Maricopa County, Arizona, USA (Figure 1A), which is characterized by urban and suburban neighborhoods surrounded by native Sonoran Desert vegetation. The regional climate is defined by extremely hot summers, warm temperatures throughout much of the year, low relative humidity (36% on average), and limited annual rainfall (183 mm on average), most of which occurs in summer and winter months.

**Figure 1.**
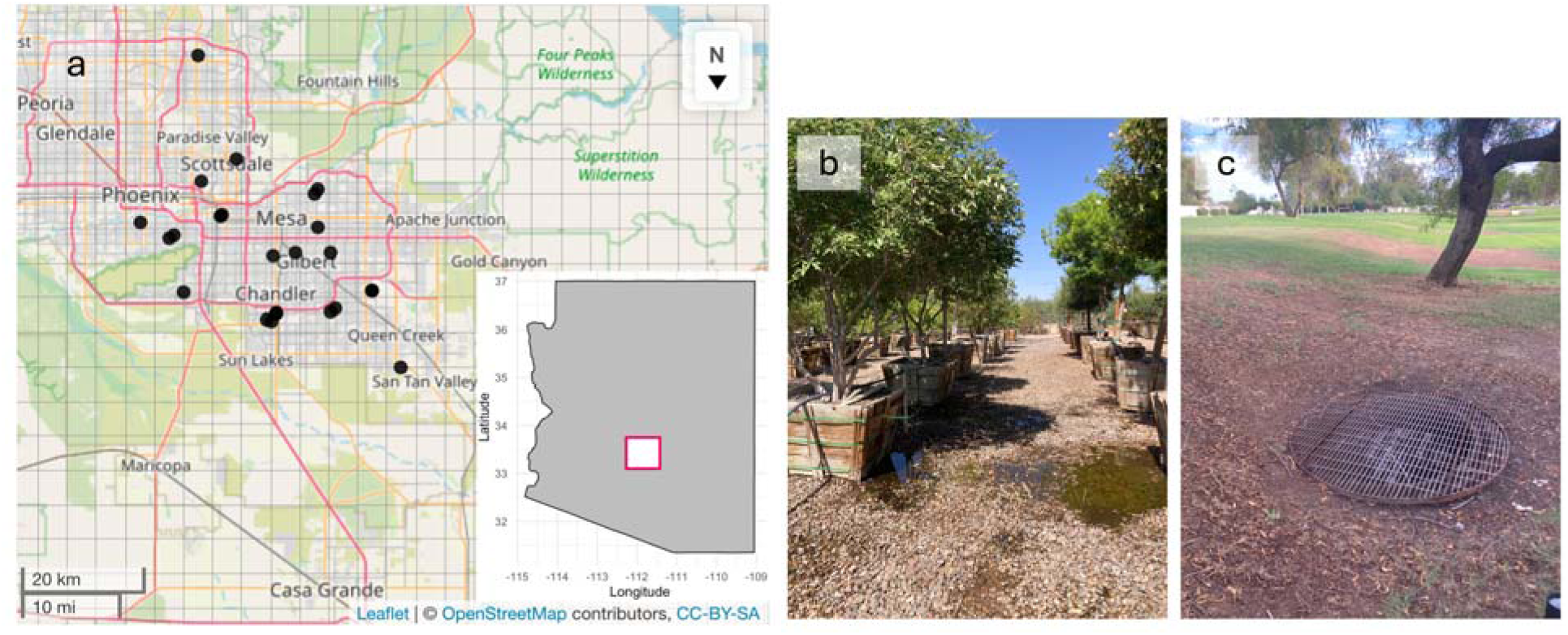
(a) Map of study sites with an inset highlighting the Phoenix metropolitan region of Maricopa County within the state of Arizona. (b) A representative photograph of a plant nursery breeding site. (c) A representative photograph of a drainage breeding site.

We identified 24 study sites across the metropolitan area covering an area of approximately 598 km². Sites were selected after inspecting hundreds of potential larval habitats across diverse land-use types, including catch basins, plant nurseries, residential parks, storm drains, grassy curb strips, culverts, roadside ditches, detention and retention ponds, and other water-retaining structures. Surveys were conducted from June 17th to August 23rd, coinciding with the peak summer heat. At plant nurseries, larvae concentrated near heavily watered trees and shrubs where shallow pools of standing water had formed (Figure 1B). In parks, larvae were mostly found in drains surrounded by large grassy areas and large catch basins (Figure 1C). We also searched permanent water bodies such as lakes and ponds but failed to detect larvae, likely due to the frequent use of larvicides and presence of aquatic predators that reduce mosquito larval survival.

At each site where mosquito larvae were detected, we sampled up to 50 individuals or as many as could be collected within 10 minutes using a standard dipper or turkey baster. Immediately following larval collection, a HOBO® temperature logger (Onset Computer Corporation, Bourne, MA, USA) was placed on the surface of the water or as close to the aquatic habitat as possible to capture microhabitat temperature profiles. The water depth in most habitats was less than a few inches deep. Collected larvae were transported to the laboratory, reared to adulthood under ambient conditions, and identified to species using standard morphological keys (Darsie and Ward 2005).

Canopy cover was quantified in situ by photographing the overhead sky at each microhabitat with an iPhone 6SE. Images were processed using ImageJ by converting photographs to binary format and analyzing the proportion of dark pixels with the “analyze particles” function to estimate fractional canopy cover.

### Temperature Data

We acquired two sources of temperature data for each larval microhabitat: on-site logger data and interpolated macroclimate data. HOBO loggers recorded ambient microhabitat temperature at 30-minute intervals (on the hour and half hour) continuously for one week following logger deployment at each site. These logger data provided fine-scale, temporal resolution measurements of local thermal regimes experienced by mosquito immature stages.

For macroclimate estimates, we obtained hourly temperature data via the Open-Meteo Historical Weather API (Zippenfenig 2023). The Open-Meteo generates 9 km gridded hourly weather variables by combining climate reanalysis models (e.g. ERA5 and ERA5-Land) and weather station data. We downloaded hourly 2 m temperature time series at the grid cell nearest each larval sampling site for the corresponding weekly periods of logger deployment.

### Statistical Analysis

We compared microhabitat temperature data with interpolated weather station records to quantify thermal buffering. Specifically, we calculated (i) the average difference in temperature between microhabitats and macroclimate estimates, (ii) the time of day when these differences were greatest, and (iii) the frequency with which microhabitat temperatures exceeded the estimated critical thermal maximum (CT_max_) of each species as reported in Mordecai et al. (2019).

To test the influence of local habitat characteristics on microclimate variation, we fit linear mixed-effects models with microhabitat temperature as the response variable. Fixed effects included macroclimate temperature estimates, percent canopy cover, and species identity, with an interaction term between weather station temperature and canopy cover to test whether shaded habitats diverged more strongly from macroclimate estimates than open habitats. Study site was included as a random intercept to account for repeated measurements. Because residual variance increased with fitted values, we modeled heteroscedasticity using an exponential variance function “varExp”. Model fitting was performed with the “nlme” package in R version 4.1.1 (Pinheiro et al. 2021).

## Results

In total, 24 larval breeding sites were identified and monitored. *Cx. quinquefasciatus* were found in the majority of sites, occupying 16 of the 24 sites, with *Cx. tarsalis* occupying the 8 remaining sites. *Cx. quinquefasciatus* were found mostly in storm drains and a few plant nurseries, while *Cx. tarsalis* were found almost exclusively in plant nurseries plus a shallow puddle in a patch of well-watered grass and flood irrigation culvert. Average temperatures in our microhabitats were 5.6 (sd = 2.6) degrees cooler than macroclimates. This consistent cooling effect has the clearest visualization at sites 19 and 11 (Figure 2), where site 19 had a dense overhead canopy (86.5%) and site 11, even though it lacked strong overhead canopy (27.5%), was shaded by surrounding vegetation; both sites were located at nurseries with shallow pooled irrigation water. The slope of the relationship between macrohabitat and microclimate temperature was 0.83, meaning that not only were microclimates cooler, but this difference intensified at warmer temperatures (Table 1, Figure 3).

**Figure 2.**
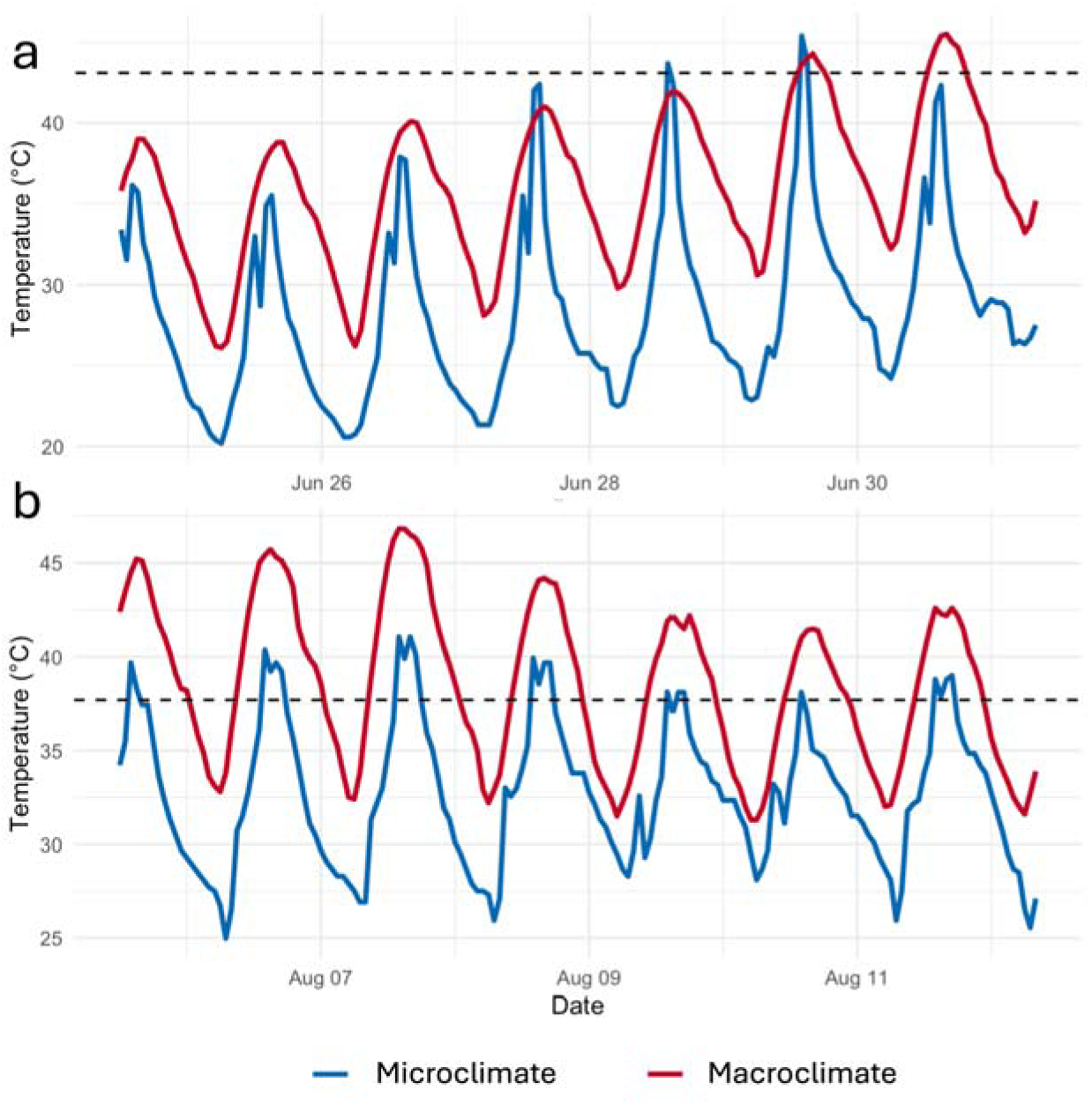
Time series of macroclimate (red) and microclimate (blue) for a breeding site of (a) *Cx tarsalis* - site 11 and (b) *Cx quinquefasciatus -* site 19.

**Figure 3.**
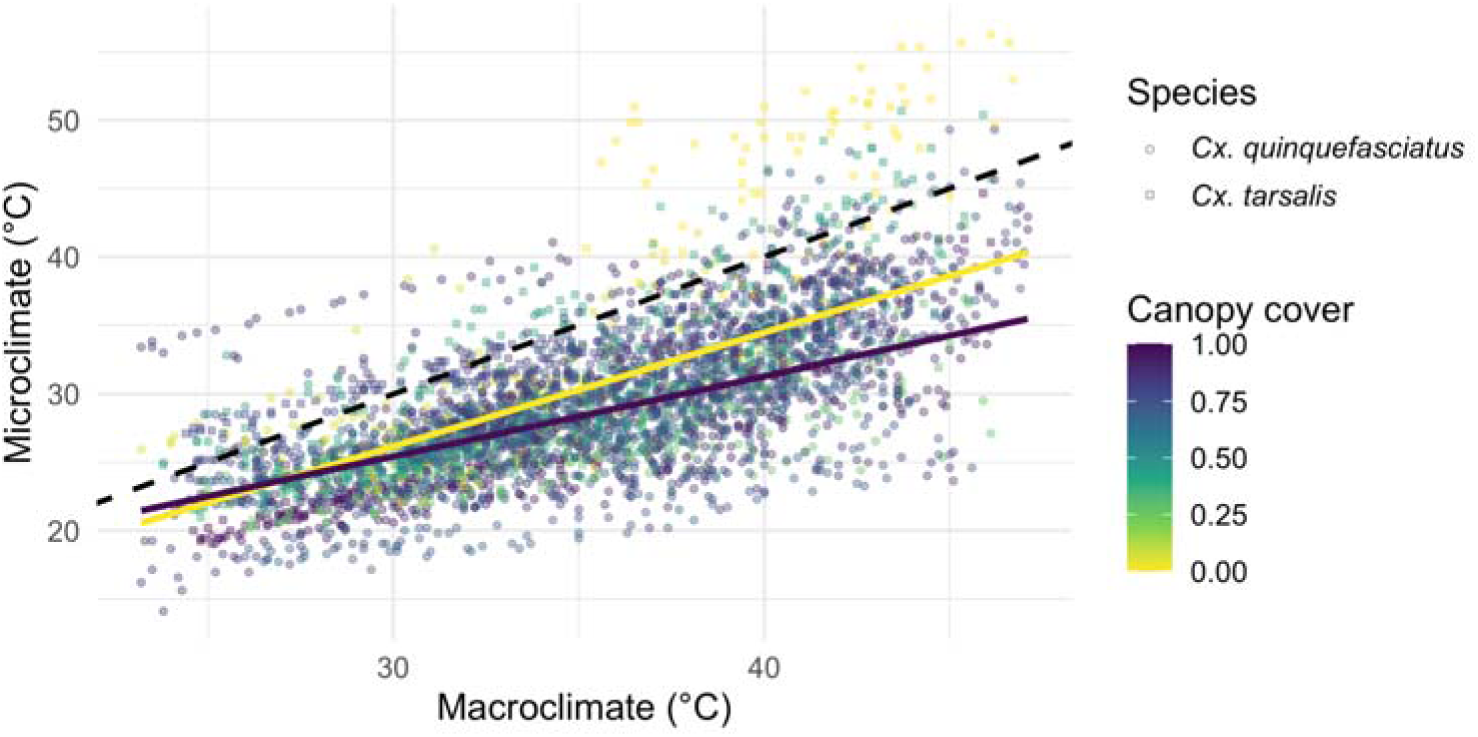
Scatterplot of macroclimate versus microclimate temperatures with a dashed 1:1 line. Points are colored by canopy cover, squares represent *Culex tarsalis* and circles represent *Culex quinquefasciatus* habitats. Lines represent predicted relationships for sites with no canopy cover in yellow and full canopy cover in blue.

**Table 1.** Summary of linear mixed-effects model predicting microhabitat temperature (°C). The macrohabitat variable was centered before the model was fit. The model included a random intercept for study site (SD = 2.47, residual SD = 0.71).

| Fixed effect | Estimate (± SE) | t-value | P-value |
| --- | --- | --- | --- |
| Intercept | 30.38 ± 1.46 | 20.75 | <0.001 |
| Macrohabitat temperature | 0.83 ± 0.03 | 30.88 | <0.001 |
| Percent canopy cover | -2.10 ± 1.92 | -1.04 | 0.31 |
| Species (*Cx. tarsalis*) | 0.86 ± 1.13 | 0.73 | 0.47 |
| Macrohabitat temperature ×<br>Percent canopy cover | -0.24 ± 0.04 | -6.49 | <0.001 |

We also tested the effects of canopy cover as a predictor of microclimate temperature. While there was not a significant effect of canopy cover on microclimate, there was a significant interactive effect between macroclimate temperature and percent canopy cover (-0.24, p < 0.001), showing that canopy can have a buffering effect, especially at higher temperatures (Figure 3).

There was also a clear distinction between time of day and the temperature differential between macroclimate and microclimate (Figure 4). The largest difference was observed in the evening between 7:00 PM to 9:00 PM, with the least difference in the morning between 5:00 AM and 7:00 AM. While the difference generally decreased during the night and increased throughout the day, there was a brief but distinct dip in the early afternoon.

**Figure 4.**
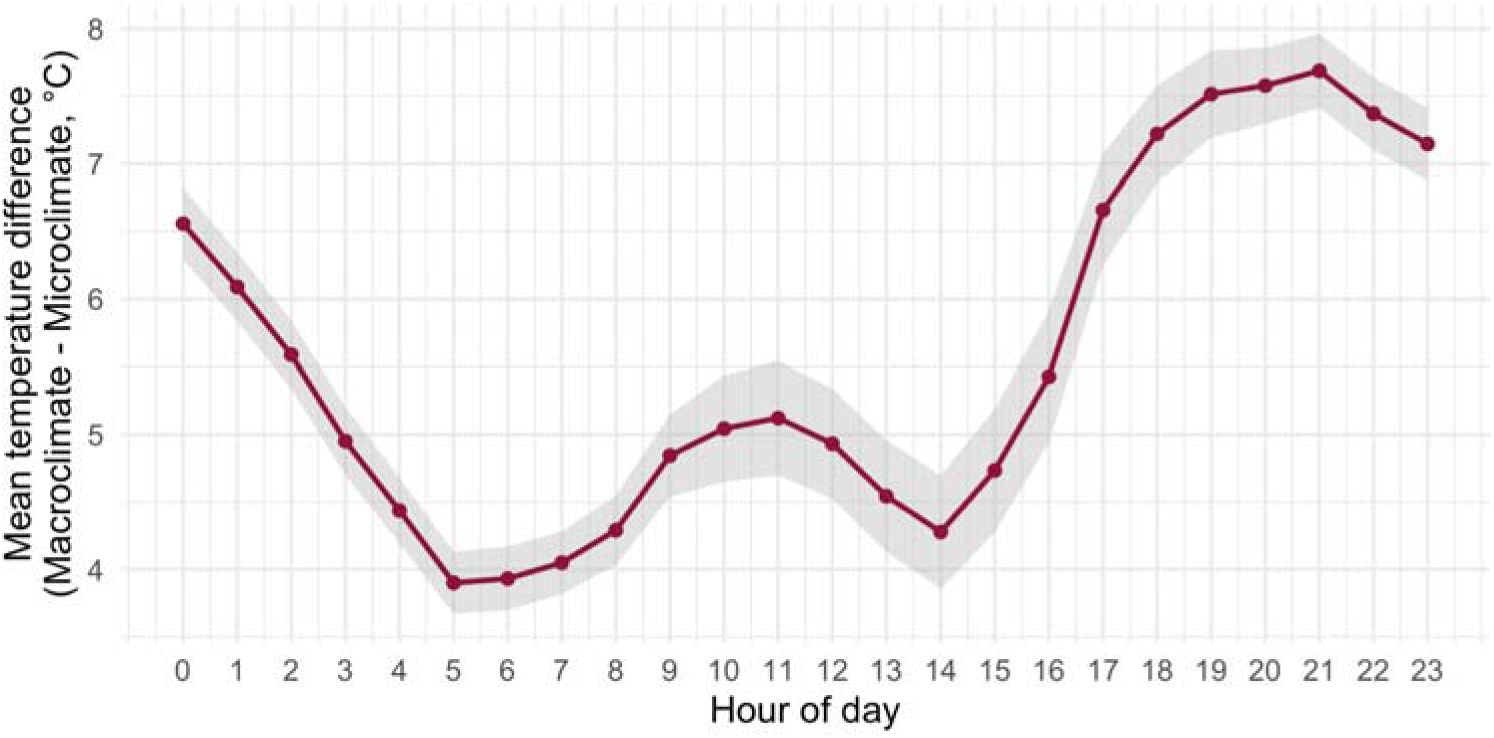
Average hourly difference in temperatures. Shaded area represents ± 1 SE.

We next compared our recorded larval habitat temperatures to the estimated CT_max_ of the species found there (Figure 5). CT_max_ for larval survival was reported by Mordecai et al (2019) as 37.7°C for *Cx. quinquefasciatus* and 43.1°C for *Cx. tarsalis*. Macroclimate data showed a significant exceedance of this thermal maxima especially for *Cx. quinquefasciatus*, with an average of over 9 hours a day exceeding CT_max_. On the other hand, microclimate exceedance was far lower, with *Cx. quinquefasciatus* CT_max_ being exceeded for only about 1 hour a day. *Cx. tarsalis* breeding sites were much less likely to exceed the CT_max_ and did so for roughly 1 hour a day as measured by both microclimate and macroclimate.

**Figure 5.**
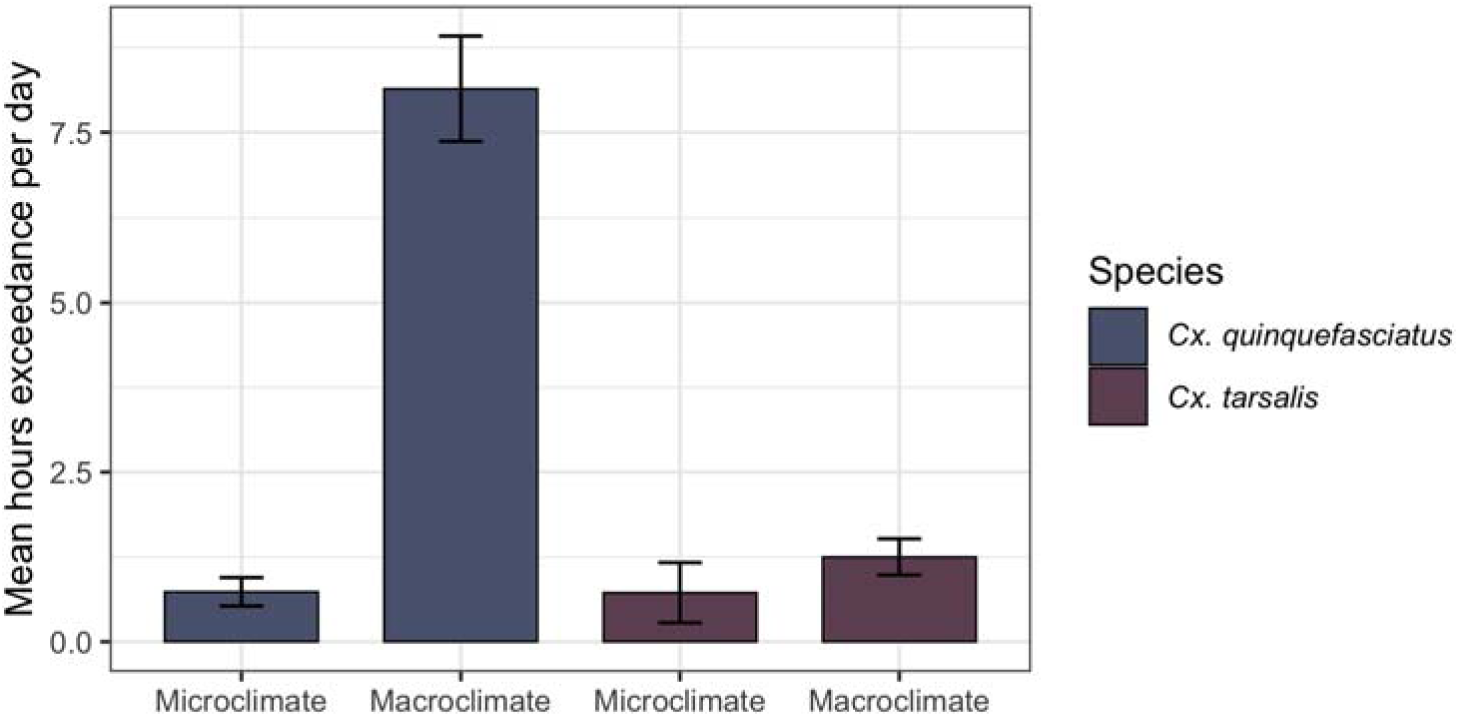
The number of hours (± 1 SE) that larval breeding sites exceeded species’ critical thermal limits.

## Discussion

Our study demonstrates that larval microhabitats of *Culex* mosquito larvae in an urban desert environment are substantially buffered from extreme heat, averaging 5.6°C cooler than macroclimate estimates. This thermal offset was most pronounced during peak heat hours (Figure 2), suggesting that local microhabitat structure provides a refuge when ambient temperatures are near or above species’ upper thermal limits. This buffering offers a necessary explanation for how small-bodied ectotherms persist in urban desert environments where ambient macroclimates frequently exceed their known upper thermal limits. For *Cx. quinquefasciatus*, macroclimate data suggested frequent exceedance of its critical thermal maximum, while microclimate records revealed exceedances of only about one hour daily. These findings indicate that gridded macroclimate datasets commonly used in mosquito ecology and disease modeling overestimate thermal stress and could therefore underestimate survival potential in hot environments.

While the general concept of microclimates mediating temperature extremes is well-documented across insect ecology, particularly in terrestrial habitats (Montejo-Kovacevich et al. 2020; Pincebourde and Woods 2020; Greiser et al. 2022) and urban habitats (Murdock et al. 2017; Wimberly et al. 2020), this buffering is especially crucial for confined aquatic ectotherms in hot, arid environments. The consistent cooling effect we observed fundamentally alters the thermal landscape for larval *Culex*. Instead of operating at or above their critical thermal maxima (CT_max_), this strong thermal refuge shifts the larval thermal regime away from lethal limits and likely closer to the optimal temperatures for development and survival (T_opt_) ( Ciota et al. 2014; Mordecai et al. 2019). This pattern parallels findings across other insects where microhabitat site selection can mitigate exposure to stressful thermal conditions (Kessler and Guerin 2008; Braem et al. 2023; Terlau et al. 2023; Vives-Ingla et al. 2023). Furthermore, for mosquitoes, living in warm but sub-lethal temperatures can accelerate disease dynamics, where warmer temperatures allow for faster viral replication, reducing the extrinsic incubation period of WNV (Reisen et al. 2006; Shocket et al. 2020; Vollans et al. 2024), thereby increasing transmission potential. Therefore, the buffering mechanism observed not only ensures mosquito survival in otherwise unsuitable conditions but may also accelerate the vector competence of the surviving population, increasing overall disease risk.

The thermal buffering observed was dynamic and strongly influenced by both ambient temperature and structural habitat features. The divergence between macroclimate and microclimate intensified at higher ambient temperatures (Figure 2), indicating maximum refuge at times when environmental stress was greatest. Our statistical model confirmed that habitat structure played an important role, showing a significant interactive effect between macroclimate temperature and percent canopy cover (Table 1). This pattern where vegetation mitigates extreme heat more strongly under high-temperature conditions has also been observed in other urban systems (Jiao et al. 2024, Ren et al. 2024). The influence of habitat structure also helps explain the distinct temporal pattern of thermal difference observed across the day (Figure 4).

The largest cooling differential occurred during the late afternoon and evening hours as well as late morning, likely reflecting the combined effects of continued structural shading from surrounding vegetation or urban infrastructure (such as storm drain walls). The brief reduction in the buffering effect during early afternoon may occur when the sun is directly overhead, reducing the effectiveness of lateral shading (Meili et al. 2021). These findings emphasize that local vegetation and urban structure create microclimatic mosaics.

Our findings have direct implications for vector population modeling and public health forecasting. Models of mosquito abundance and pathogen transmission often rely on interpolated or reanalysis climate data (e.g. ERA 5 Land at a 9km resolution, PRISM at a 4km resolution (Daly et al. 2008; Muñoz-Sabater 2021)) or nearby weather stations to approximate the thermal experience of mosquitoes (Soh and Aik 2021; Ferraccioli et al. 2023; Gorris et al. 2023; Kessinger et al. 2025). Similar assumptions are common in population, phenology, and pest forecasting models across insect systems, where coarse climate data are often used as proxies for experienced temperatures (Hodgson et al. 2011, Rebaudo et al. 2026). However, our results suggest that this assumption may lead to systematic errors, because macroclimate products smooth over the fine scale thermal mosaics created by shading, containment, water depth, and built structure. In warm cities where these features are common, such averaging produces thermal environments that are hotter, more exposed, and more variable than those actually experienced by mosquito larvae. Recent work shows that explicitly modeling the microclimatic temperature in storm drains can markedly change predictions of mosquito population dynamics (Erraguntla et al. 2021).

There have been important advances in generating high resolution climate models that incorporate shading and radiation, such as SOLWEIG (Lindberg et al. 2008), yet their spatial extent, data requirements, and computational cost limit their ability to provide city wide inputs for mosquito models. Some climate data products attempt to include microclimate adjustments by modifying temperature inputs according to land cover, canopy fraction, or water body presence (Zellweger et al. 2019). Mechanistic thermal ecology tools such as Microclim and NicheMapR aim to estimate organism level temperatures by modeling heat exchange processes. These approaches capture elements of microclimate variation but still lack direct measurements of small aquatic habitats like the ones studied here. A continuing challenge is to develop scalable methods that translate coarse climate data into biologically relevant thermal conditions without requiring extensive site specific measurements (Potter et al. 2013).

Our sampling period was limited to summer months to sample microclimate dynamics under extreme heat. While this highlights survival strategies during peak thermal stress, it does overlook seasonal variation, like whether these buffering effects are as prominent during cooler seasons, where previous studies have found winter buffering can also be substantial (Trewin et al. 2019, Romiti et al. 2025). The summer also lacked any significant rainfall, with the Flood Control District of Maricopa County reporting approximately 3-5 cm falling during our study period. Thus our data does not account for the impact of precipitation, which can reshape microclimates by increasing humidity, altering thermal stability, and generating additional transient wet habitats (Ruiz et al. 2010, Brown et al. 2023). Beyond extending temporal observations, further data collection should focus on gathering more diverse habitat types of *Culex* mosquitoes, including those of adults, as the majority of our data was collected from plant nurseries or storm drains which don’t account for other potential buffering strategies in habitats such as vegetation, river beds, and basements. Doing so leads to the potential to develop scalable correction factors to translate macroclimate data into microclimate estimates. As our priority in this study was to monitor active breeding sites, there was no focus on capturing the extent of data necessary for a robust macroclimate to microclimate correction. Further climate data, landscape features, habitat type, and vegetation needs to be collected in areas with and without mosquito presence if a corrective model is to be created.

In conclusion, our results highlight that mosquito larval habitats experience microclimates substantially cooler than ambient macroclimate conditions, providing thermal refugia that likely sustain populations in otherwise inhospitable urban deserts. Accounting for these fine-scale temperature differences will improve the realism of insect ecology and transmission models, but incorporating them will require targeted microclimate measurements or validated correction approaches that account for vegetation, structural shading, water body properties, and temporal variation.

## Acknowledgements

This material is based upon work supported by the National Science Foundation under grant number(s) DEB-2224662, Central Arizona-Phoenix Long-Term Ecological Research Program (CAP LTER). USDA NIFA award 2024 77040 43178 supported AV. We thank the Maricopa County Vector Control Division for sharing their expertise on larval mosquito habitats.

## Author Contributions

https://academic.oup.com/ee/pages/Manuscript_Preparation

KL - Conceptualization, Investigation, Methodology, Supervision, Writing – original draft

CM - Conceptualization, Formal Analysis, Investigation, Visualization, Writing – original draft

AV - Data Curation, Investigation, Writing – original draft

